# Signaling Pathway Activities Improve Prognosis for Breast Cancer

**DOI:** 10.1101/132357

**Authors:** Yunlong Jiao, Marta R. Hidalgo, Cankut Çubuk, Alicia Amadoz, José Carbonell-Caballero, Jean-Philippe Vert, Joaquín Dopazo

## Abstract

With the advent of high-throughput technologies for genome-wide expression profiling, a large number of methods have been proposed to discover gene-based signatures as biomarkers to guide cancer prognosis. However, it is often difficult to interpret the list of genes in a prognostic signature regarding the underlying biological processes responsible for disease progression or therapeutic response. A particularly interesting alternative to gene-based biomarkers is mechanistic biomarkers, derived from signaling pathway activities, which are known to play a key role in cancer progression and thus provide more informative insights into cellular functions involved in cancer mechanism. In this study, we demonstrate that pathway-level features, such as the activity of signaling circuits, outperform conventional gene-level features in prediction performance in breast cancer prognosis. We also show that the proposed classification scheme can even suggest, in addition to relevant signaling circuits related to disease outcome, a list of genes that do not code for signaling proteins whose contribution to cancer prognosis potentially supplements the mechanisms detected by pathway analysis.

## Introduction

Over the past decades, many efforts have been addressed to the identification of gene-based signatures to predict patient prognosis using gene expression data (Paik, et al., 2004; Sotiriou and Pusztai, 2009; van ′t Veer, et al., 2002; Wang, et al., 2005). Despite the success of its use, gene expression signatures have not been exempt of problems (Ein-Dor, et al., 2006; Iwamoto and Pusztai, 2010). Specifically, one major drawback of multi-gene biomarkers is that they often lack proper interpretation in terms of mechanistic link to the fundamental cell processes responsible for disease progression or therapeutic response (Dopazo, 2010; van′t Veer and Bernards, 2008). Actually, it is increasingly recognized that complex traits, such as disease or drug response, are better understood as alterations in the operation of functional modules caused by different combinations of gene perturbations (Barabasi, et al., 2011; Barabasi and Oltvai, 2004; Oti and Brunner, 2007). To address this inherent complexity different methodologies have tried to exploit several functional module conceptual representations, such as protein interaction networks or pathways, to interpret gene expression data within a systems biology context (Barabasi, et al., 2011; Fryburg, et al., 2014; Hood, 2013; Vidal, et al., 2011). Actually, it has recently been shown that the pathway-level representation generates clinically relevant stratifications and outcome predictors for glioblastoma and colorectal cancer (Drier, et al., 2013) and also breast cancer (Livshits, et al., 2015). Moreover, mathematical models of the activity of a pathway have demonstrated a significantly better association to poor prognosis in neuroblastoma patients than the activity of their constituent genes, including *MICN*, a conventional biomarker (Fey, et al., 2015). This observation has recently been extended to other cancers (Hidalgo, et al., 2017) and to the prediction of drug effects (Amadoz, et al., 2015).

Given that the inferred activity of the pathway should be closely related to its cellular mechanism for disease progression, its use to guide cancer prognosis seems promising. Recently, a number of pathway activity inference methods have been proposed (Hidalgo, et al., 2017; Jacob, et al., 2012; Li, et al., 2015; Martini, et al., 2013). Here, we use the canonical circuit activity analysis method, which has demonstrated to have a superior performance (Hidalgo, et al., 2017) finding significant associations of specific circuit activities, directly responsible for triggering the prominent cancer hallmarks (Hanahan and Weinberg, 2011), to patient survival. This method recodes gene expression values into measurements of signaling circuit activities that ultimately account for cell responses to specific stimuli. Such activity values can be considered multigenic mechanistic biomarkers that can be used as features for cancer prognosis.

We demonstrate that the activity of signaling circuits yields comparable or even better prediction in breast cancer prognosis than the expression of individual genes, while detected mechanistic biomarkers enjoy the compelling advantage of readily available interpretation in terms of the corresponding cellular functions they trigger. Moreover, we show that the proposed prediction scheme can even suggest, in addition to interesting signaling circuits related to disease outcome, a list of prognostic genes that do not code for signaling proteins whose contribution to cancer prognosis potentially supplements the mechanism included in the pathways modeled.

## Methods

### Data source and processing

The breast cancer gene expression and survival data used here was downloaded from The Cancer Genome Atlas (TCGA), release No. 20 of the International Cancer Genome Consortium (ICGC) data portal (https://dcc.icgc.org/releases/release_20/Projects/BRCA-US). This dataset provides the RNA-seq counts of 18,708 genes for 879 tumor samples, in which we also have records of the vital status of corresponding donors, namely the overall survival outcome of the cancer patients being alive or deceased at the end of clinical treatment (Table 1). Since TCGA cancer data are from different origins and underwent different management processes, non-biological experimental variations, commonly known as batch effect, associated to Genome Characterization Center (GCC) and plate ID must be removed from the RNA-seq data. The COMBAT method (Johnson, et al., 2007) was used for this purpose. We then applied the trimmed mean of M-values normalization method (TMM) method (Robinson and Oshlack, 2010) for data normalization which is essential in applying the CCAA method. The resulting normalized values were finally entered to the pathway analysis method.

**Table 1.**
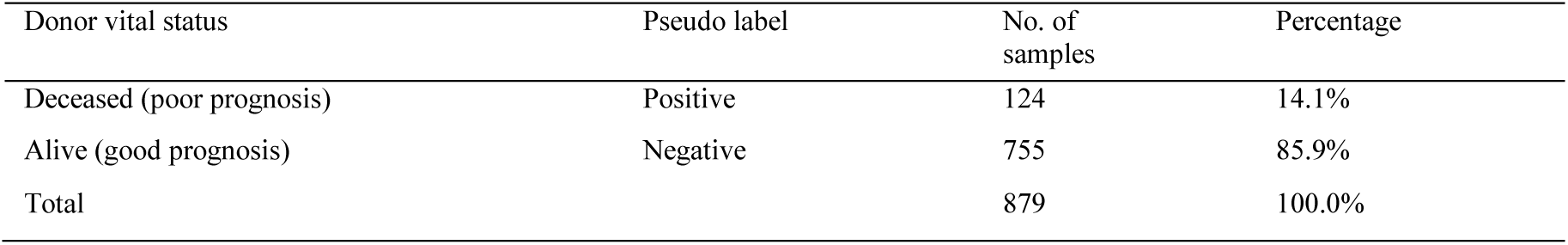
Summary of survival outcome of the breast cancer patients in the TCGA dataset.

| Donor vital status | Pseudo label | No. of samples | Percentage |
| --- | --- | --- | --- |
| Deceased (poor prognosis) | Positive | 124 | 14.1% |
| Alive (good prognosis) | Negative | 755 | 85.9% |
| Total |  | 879 | 100.0% |

A total of 60 KEGG pathways (Supplementary Table 1) were downloaded from the KEGG repository (Kanehisa, et al., 2012), including 2,212 gene products that participate in 3,379 nodes.

### Modeling framework for signaling pathways

We applied the canonical circuit activity analysis method (Hidalgo, et al., 2017), as implemented in the *hipathia* R package available at https://github.com/babelomics/hipathia, in pursuit of modeling signaling activity. Within the modeling context, a *circuit* is defined as all possible routes the signal can traverse to be transmitted from a particular input node to a particular output node (see Supplementary Figure 1A). A total of 6,101 *circuits* are identified and modeled in this study. The transmission of the signal depends on the integrity of the chain of nodes that connect the receptor to the effector and briefly, it is estimated as follows. The presence of the mRNA (the normalized RNA-seq counts rescaled between 0 and 1) is taken as a proxy for the presence of the corresponding protein in each pathway node (Bhardwaj and Lu, 2005; Efroni, et al., 2007; Montaner, et al., 2009; Sebastian-Leon, et al., 2014). Then, the degree of integrity of the *circuit* is estimated by modeling the signal flow across it. Specifically, the input node (receptor) is initialized by an incoming signal of intensity value of 1, and then for each node *n* of the *circuit*, the signal value *sn* is updated by the following rule:

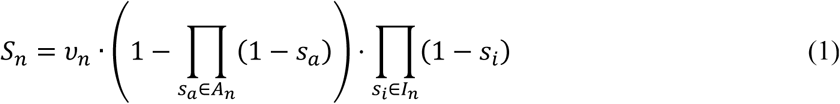

where *An* denotes the set of signals arriving to the node from activation edges, *In* denotes the set of signals arriving to the node from inhibition edges, and *vn* is the (normalized) value of the current node *n*.

**Fig. 1.**
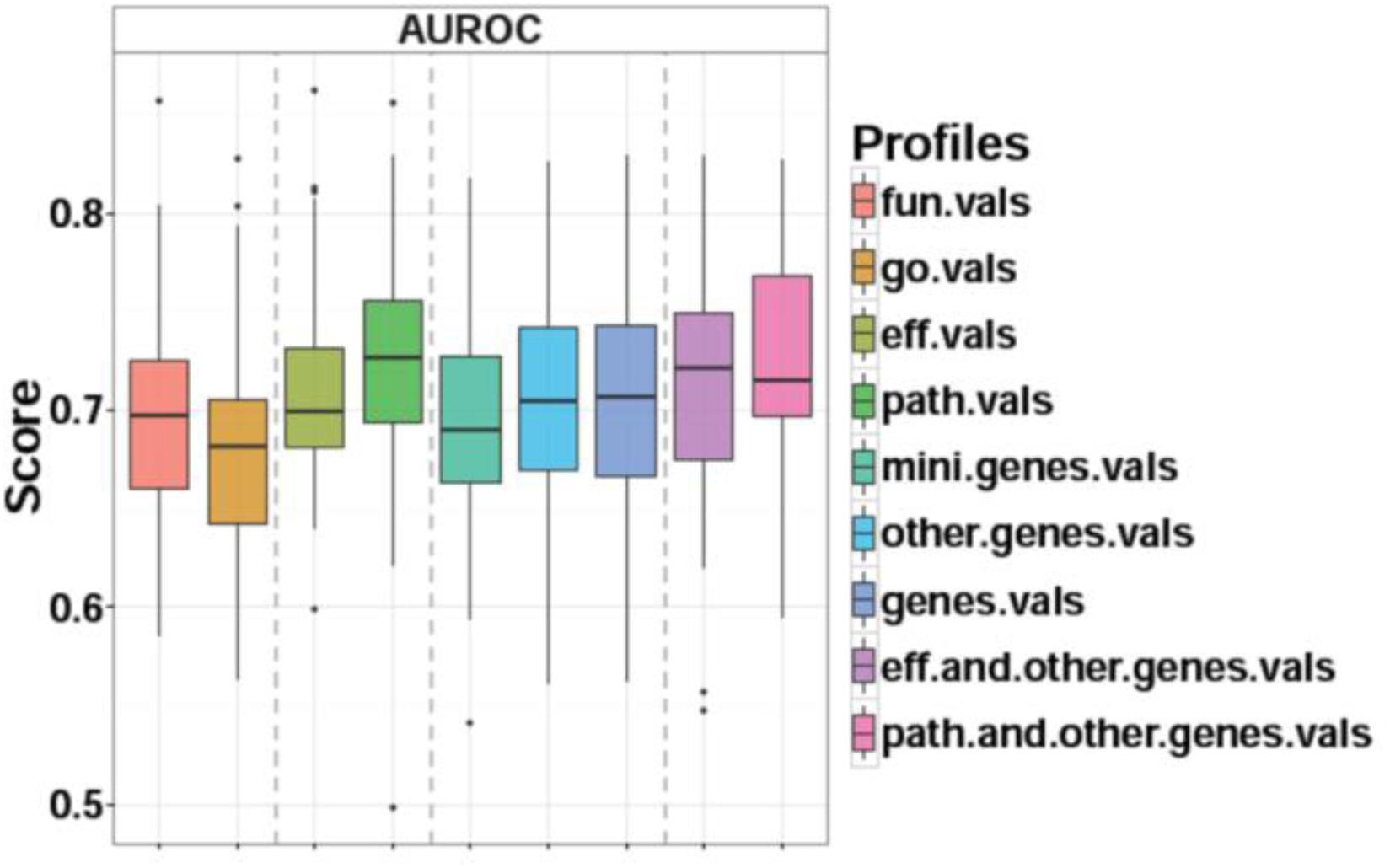
The AUROC performance of using different types of profiles as predictive features to classify survival outcome for breast cancer patients. Boxplot represents the variance of the performance on 50 cross-validation splits. Dotted vertical lines separate profiles by the underlying analysis levels.

Finally, the activity value for the *circuit* is defined by the signal intensity transmitted through the last (effector) protein of the circuit which quantifies the cell function ultimately activated by the *circuit*.

Since output nodes at the end of *circuits* are the ultimate triggers of specific cellular actions, an *effector circuit* is defined from a functional viewpoint as a higher-level signaling entity that composes all *circuits* ending at the same output node. When applied to an effector circuit, the method returns the joint intensity of the signal arriving to the corresponding effector node (see Supplementary Figure 1B). Furthermore, the known functions triggered by each effector protein in cell can be derived from their functional annotations. Here we use UniProt (UniProt_Consortium, 2015) and Gene Ontology (Ashburner, et al., 2000) (GO) annotations.

Finally, inferred signaling activity values of those effector circuits ending at proteins with the same annotated functions are averaged to quantify the activity of the function realized in cell. This way we obtain estimated activity values directly connected to a list of cellular functions (Supplementary Figure 1C).

Supplementary Figure 1 depicts the different levels of abstraction from *circuits*, to *effector circuits* and finally functions. Eventually, a subset of curated functions can be used for a specific scenario in which the relevant functions are known. Here we use cancer hallmarks (Hanahan and Weinberg, 2011).

### Cancer prognosis with inferred signaling pathway activity

In this study, we are interested in evaluating the prognostic power of pathway-level mechanistic features and gene-based features alone and in combination. Using the *hipathia* method we recoded the list of gene expression values of each tumor sample into the corresponding lists of signaling activity values for the three levels of abstraction: *circuits, effector circuits* and functions, as described in UniProt and GO annotations. Therefore for each tumor sample we end up with a profile of gene expression, a profile of *circuit* signaling activity, a profile of *effector circuit* signaling activity, a profile of UniProt-based cellular function activity and a profile of GO-based cellular function activity. These profiles are sample-specific profiles that can be straightforwardly used as prognostic features using any classification algorithm. Note that pathway-level profiles are derived with no regard to any information provided by the genes whose products do not participate in cell signaling, and the prognostic power of pathway-level profiles may thus be limited by the coverage of genes in known biological pathways. In order to understand the relative contribution to the pathway-level profiles and gene-level profiles to the accurate separation between good vs poor prognosis, we devised four artificial profiles: *path-gene* expression profile containing only genes that are involved in the KEGG signaling pathways, *other-gene* expression profile containing only genes that are absent from the KEGG pathways, a combined profile consisting of signaling activity of effector circuits and expression of other-genes, and a combined profile consisting of signaling activity of *circuits* and expression of other-genes. Thus, we use a total of 9 types of profiles (detailed in Table 2) From the viewpoint of machine learning, this study is formulated as a typical binary classification problem where we determine a positive or negative pseudo label for each sample. Based on the data available in this study (Table 1) we perform a 5-fold cross-validation repeated 10 times on the dataset and report the mean performance over the 5 *×* 10 = 50 splits to assess the prognostic power for each type of profile. The performance is evaluated by the Area Under the ROC Curve (AUROC) criteria (Sing, et al., 2005). Note that usually a classifier returns a continuous prediction between 0 and 1 for each sample denoting the probability of that sample being in the positive class rather than in the negative class, and then assigns either label to the sample according to some cutoff value thresholding the prediction. In fact, AUROC is a cutoff-free score that measures the probability that the classifier will score a randomly drawn positive sample higher than a randomly drawn negative sample.

**Table 2.** Summary of different types of profiles used as predictive features for breast cancer prognosis in this study.

| Alias | Profile type | Number features | Analysis level |
| --- | --- | --- | --- |
| fun.vals | UniProt-based functions | 81 | Circuit- |
| go.vals | GO-based functions | 370 | Circuit- |
| eff.vals | Effector circuits | 1,038 | Circuit- |
| path.vals | Circuits | 6,101 | Circuit- |
| path.genes.vals | Pathway-genes | 2,212 | Gene |
| other.genes.vals | Other-genes | 16,496 | Gene |
| genes.vals | All genes | 18,708 | Genes |
| eff.and.other.- genes.vals | Effector and other-genes | 17,534 | Circuit + Gene |
| path.and.other.- genes.vals | Circuits and other-genes | 22,597 | Circuit + Gene |

In this study we consider 12 classification algorithms as candidate classifiers, most of which are state-of-the-art (Table 3). When we assess the prognosis performance for a specific type of profile on a specific (external) cross-validation split of the data, we perform an internal 5-fold cross-validation on the training set to determine which classifier returns the highest cross-validated performance and the best classifier is then used on the test set to obtain the performance score. This procedure guarantees that the performance on each (external) cross-validation split is evaluated impartially for each profile with its best suited algorithm.

**Table 3.** The 12 classifiers considered in this study to classify prognosis for breast tumor samples. Note that majority voting classifier serves as a baseline negative-control model which outputs a constant label for any test sample by the dominant class in the training set..

| Alias | Classifier | Reference |
| --- | --- | --- |
| LDA | Linear discriminant analysis | (Ripley, 2007; Venables and Ripley, 2002) |
| LogitLasso | L1-regularized logistic regression | (Friedman, et al., 2010) |
| LinearSVM | Support Vector Machine with linear kernel | (Chang and Lin, 2011) |
| RadialSVM | Support Vector Machine with Gaussian RBF kernel | (Chang and Lin, 2011) |
| KendallSVM | Support Vector Machine with Kendall kernel | (Jiao and Vert, 2016; Zeileis, et al., 2004) |
| KNN | k-nearest neighbor classifier | (Ripley, 2007; Venables and Ripley, 2002) |
| NB | Naive Bayes classifier | (Ripley, 2007) |
| GBM | Gradient Boosting Machine | (Friedman, 2001) |
| RF | Random Forest | (Breiman, 2001; Liaw and Wiener, 2002) |
| SparseSVM | L1-regularized L2-loss Support Vector Machine | (Fan, et al., 2008) |
| PAM | Nearest shrunken centroid classifier | (Tibshirani, et al., 2002) |
| Constant | Majority voting classifier | — |

## Results

### 3.1 Signaling pathway activity leads to improved prognosis for breast tumor samples

The performance of using different types of profiles (Table 2) as predictive features to classify survival outcome for breast cancer patients is shown in Figure 1 Under either criterion of AUROC to evaluate the classification performance, we observe that the activity values of signaling *circuits,* denoted by *path.vals*, yield the best performance overall. In particular, they outperform the profiles based solely on gene expression values, denoted by *path.genes.vals, other.genes.vals* and *genes.vals*. In other words, we are able to integrate the expression values of *path-genes* into the *a priori* knowledge of cell signaling to obtain pathway-level features that achieve improved prognosis. Interestingly, these pathway-level features relate to biological processes and cellular functions *per se*. Although the pathway-level features are derived from the expression of *path-genes* and thus agnostic to the expression of *other-genes*, the inclusion of *other-genes* to the signaling circuits activity values, denoted by *eff.and.other.genes.vals* and *path.and.other.genes.vals* profiles, does not significantly improve the performance (no significant differences after applying a two-sided t-test comparing differences between the cross-validation AUROC scores obtained by each pair of profiles, and adjusted for multiple testing (Benjamini and Hochberg, 1995), see Table 4).

**Table 4.** FDR-adjusted p-values comparing the corresponding classification scores of feature s in columns versus features in files over 50 cross-validation splits. See Figure 1 for the performance values of each feature. Significant values are in boldface and marked with an asterisk..

|  | 1 | 2 | 3 | 4 | 5 | 6 | 7 | 8 |
| --- | --- | --- | --- | --- | --- | --- | --- | --- |
| 1 *fun.vals* | 0.1467 | 0.1306 | **0.0119\*** | 0.8267 | 0.1908 | 0.1908 | 0.0815 | **0.002\*** |
| 2 *go.vals* |  | **0.0024\*** | **0.0004\*** | 0.1557 | **0.0120\*** | **0.0119\*** | **0.0046\*** | **<0.0001\*** |
| 3 *eff.vals* |  |  | 0.0815 | **0.0255\*** | 0.8743 | 0.8743 | 0.6422 | **0.0235\*** |
| 4 *path.vals* |  |  |  | **0.0046\*** | 0.1408 | 0.1394 | 0.3702 | 0.6422 |
| 5 *path.genes* |  |  |  |  | **0.0255\*** | **0.0255\*** | **0.0046\*** | **0.0002\*** |
| 6 *other.genes* |  |  |  |  |  | 0.9483 | 0.2167 | **0.0039\*** |
| 7 *genes* |  |  |  |  |  |  | 0.2 | **0.0032\*** |
| 8 *eff.and.other.genes* |  |  |  |  |  |  |  | **0.0473\*** |

When comparing the prognostic power between pathway-level and gene-level profiles, we have also derived cellular function activity profiles, denoted by *fun.vals* and *go.vals* (Table 2), and observed that the performance of these profiles are slightly worse than other pathway-level profiles (Fig. 1). This is probably due to the excessively simplistic procedure that basically averages the signaling activity values of effector circuits ending at proteins with the same annotated keywords according to Uniprot or GO (Hidalgo, et al., 2017), annotations that can be incomplete and ambiguous to some extent.

Table 5 summarizes the best-performing classifiers for each type of prognostic profile in the sense that they are most frequently selected by internal cross-validation. Notably, it evidences that Support Vector Machines with various kernels are recurrently selected as the competent classifier in breast cancer prognosis that suits well for both gene-level and pathway-level features.

**Table 5.**
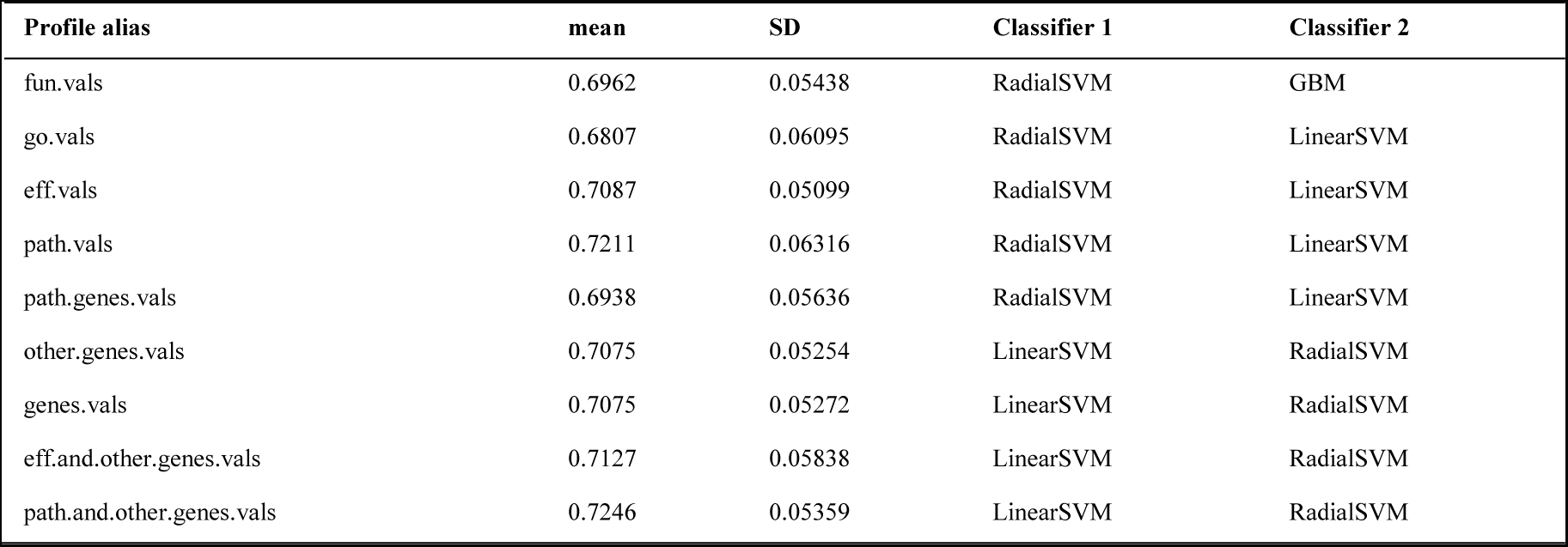
Top two most frequently selected classifiers by internal cross-validation for each type of prognostic profile in classifying breast cancer prognosis evaluated by AUROC.

| Profile alias | mean | SD | Classifier 1 | Classifier 2 |
| --- | --- | --- | --- | --- |
| fun.vals | 0.6962 | 0.05438 | RadialSVM | GBM |
| go.vals | 0.6807 | 0.06095 | RadialSVM | LinearSVM |
| eff.vals | 0.7087 | 0.05099 | RadialSVM | LinearSVM |
| path.vals | 0.7211 | 0.06316 | RadialSVM | LinearSVM |
| path.genes.vals | 0.6938 | 0.05636 | RadialSVM | LinearSVM |
| other.genes.vals | 0.7075 | 0.05254 | LinearSVM | RadialSVM |
| genes.vals | 0.7075 | 0.05272 | LinearSVM | RadialSVM |
| eff.and.other.genes.vals | 0.7127 | 0.05838 | LinearSVM | RadialSVM |
| path.and.other.genes.vals | 0.7246 | 0.05359 | LinearSVM | RadialSVM |

### 3.2 Signaling circuits selected as features relevant for cancer prognostic account for cancer hallmarks

From the clinical standpoint of cancer prognosis, we are interested in identifying a small set of biomarkers that can guide decision making in cancer prognosis. As our analysis is made at the level of pathways, we would like to detect a few signaling *circuits* whose activity, and thus the underlying cell functionality, has a significant impact on discriminating the prognosis classes of cancer patients. We opted for the Random Forest classifier to perform this analysis, since it simultaneously predicts the survival outcome of tumor samples and scores the importance of each feature that is ultimately used in the prediction. We focus on the feature importance measure returned by fitting a Random Forest which accounts for the mean decrease in classification performance if we randomly permute the data of the corresponding feature. Table 6 lists the five top-scored signaling *circuits* by fitting Random Forests with the profiles of *circuit* activities (denoted by *path.vals*). The role played by each signaling circuit in cancer progression can be inferred from the underlying cellular functions (taken from GO annotations) triggered by the last (effector) protein on the circuit. Thus, the first *circuit*, belonging to the *HIF-1 signaling pathway*, starts with the *TLR4* receptor, which is known to be related to progression of several cancers (breast, ovarian, prostate and head and neck) via Lipopolysaccharide Stimulation (Yang, et al., 2014) and ends in the *EDN1* effector, an hypoxia-inducible factor that mediates cancer progression (Semenza, 2012). Another relevant *circuit* belongs to the *NF-kappa B signaling pathway* and has the *IL1B* protein as receptor and the *CXCL2* as effector. Polymorphisms in the receptor have been linked to several cancers in different populations (El-Omar, et al., 2000; Lu, et al., 2005) and it has been demonstrated the role of *CXCL2* in tumor growth and angiogenesis (Keane, et al., 2004). Similarly, polymorphisms in the *LEP* protein, the receptor of another *circuit* in the *Adipocytokine signaling pathway*, have been linked to cancer (Cleveland, et al., 2010), and its effector, the tyrosine phosphatase Shp2 (*PTPN11*), contributes to the pathogenesis of many cancers and other human diseases (Chan, et al., 2008). The Cell cycle signaling pathway contains another relevant *circuit* whose receptor *TTK* transmits the signal until the cohesin complex. This four proteins complex is essential for chromosome segregation and DNA repair and mutations in its component genes have recently been identified in several types of tumors (Losada, 2014). Finally, the fifth most relevant *circuit*, belonging to the *Tight junction pathway,* contains the *AKT3* serine/threonine kinase with a known role in tumorigenesis (Testa and Bellacosa, 2001), is signaled by the receptor *ACTN4*, a protein which has been related to cell invasion and metastasis (Honda, 2015). Supplementary Table 2 shows an expanded list of top-scored 50 *circuits*.

**Table 6.** Top five *circuits* with the highest feature importance measure by fitting Random Forests with *path.vals* in classifying breast cancer prognosis.

| Pathway name | Receptor Gene(s) | Effector Gene(s) | Effector protein function |
| --- | --- | --- | --- |
| HIF-1 | *TLR4* | *EDN1* | Growth/survival factor in cancer |
| NF-kappa B | *IL1B* | *CXCL2* | Inflammatory response and angiogenesis |
| Adipocytokine | *LEP* | *PTPN11* | Protein phosphatase |
| Cell cycle | *TTK* | Cohesin complex<br>(*SMC1B*, *SMC3*,<br>*STAG1*, *RAD21*) | Chromosome segregation and DNA repair |
| Tight junction | *ACTN4*, *MAGI3* | *AKT3* | Cell invasion and metastasis |

Table 7 lists the top-scored *effector circuits* by fitting Random Forests with the profiles of *effector circuit* activities (denoted by *eff.vals*). Although the cohesion complex effector is again selected, the effector circuit level analysis provided a slightly different perspective of relevant aspects of signaling in cancer patient survival. Thus, two *effector circuits* with effector proteins *LEPR* and *PPARα*, from the *AMPK* and the *Adipocytokine* signaling pathways, respectively, are activators of the fatty acid metabolism. Two more effector pathways ending in the Interleukin 6 (IL6), related to inflammatory processes and immune response in the *Toll-like receptor pathway*, seem more likely to be involved in blocking the cell differentiation through the *Pathways in cancer* (KEGG id hsa05200). Actually, it has been described that *IL6* blocks apoptosis in cells during the inflammatory process, keeping them alive in toxic environments, but the same process protects cells from apoptosis and chemotherapeutic drugs during neoplastic growth (Hodge, et al., 2005). Supplementary Table 3 shows an expanded list of top-scored 50 *effector circuits*. Beyond the top scored *signaling circuits* (Table 6) and *effector circuits* (Table 7), other relevant circuits are listed in Supplementary Tables 2 and 3. Although an exhaustive list of the consequences that processes differentially activated can have in tumorigenesis is beyond the scope of this work, it is worth noticing that cancer a hallmark such as apoptosis inhibition is represented by inhibition of *signaling circuits IL6-BCL2* in the *HIF-1 signaling pathway* and IL10-BNIP3 in the *FoxO signaling pathway* (eighth and ninth in Supplementary Table 2, respectively), both containing the protein *STAT3*, known to mediate apoptosis inhibition in breast cancer (Gritsko, et al., 2006) (see Figure 2). The graphic representation of the complete *effector circuits* containing the two *signaling circuits* in the *HIF-1 signaling pathway* and the *FoxO signaling pathway* has been obtained with the hipathia web tool (Hidalgo, et al., 2017), using the *path.genes.vals* gene expression profiles (that are converted to *eff.vals* and *path.vals* profiles by the program).

**Table 7.** Top five *effector circuits* with the highest feature importance measure by fitting Random Forests with *eff.vals* in classifying breast cancer prognosis.

| Pathway name | Effector gene | Effector protein function |
| --- | --- | --- |
| AMPK | *LEPR* | Regulation of fatty acid metabolism |
| Adipocytokine | *PPARα* | Peroxisome proliferation and fatty acid metabolism |
| Pathways in cancer | *IL6* | Blockage of differentiation, Anti-apoptosis |
| Cell cycle | Cohesin complex<br>(*SMC1B*, *SMC3*,<br>*STAG1*, *RAD21*) | Chromosome segregation and DNA repair |
| Toll-like receptor | *IL6* | Inflammation, Immune response, Anti-apoptosis |

**Fig. 2.**
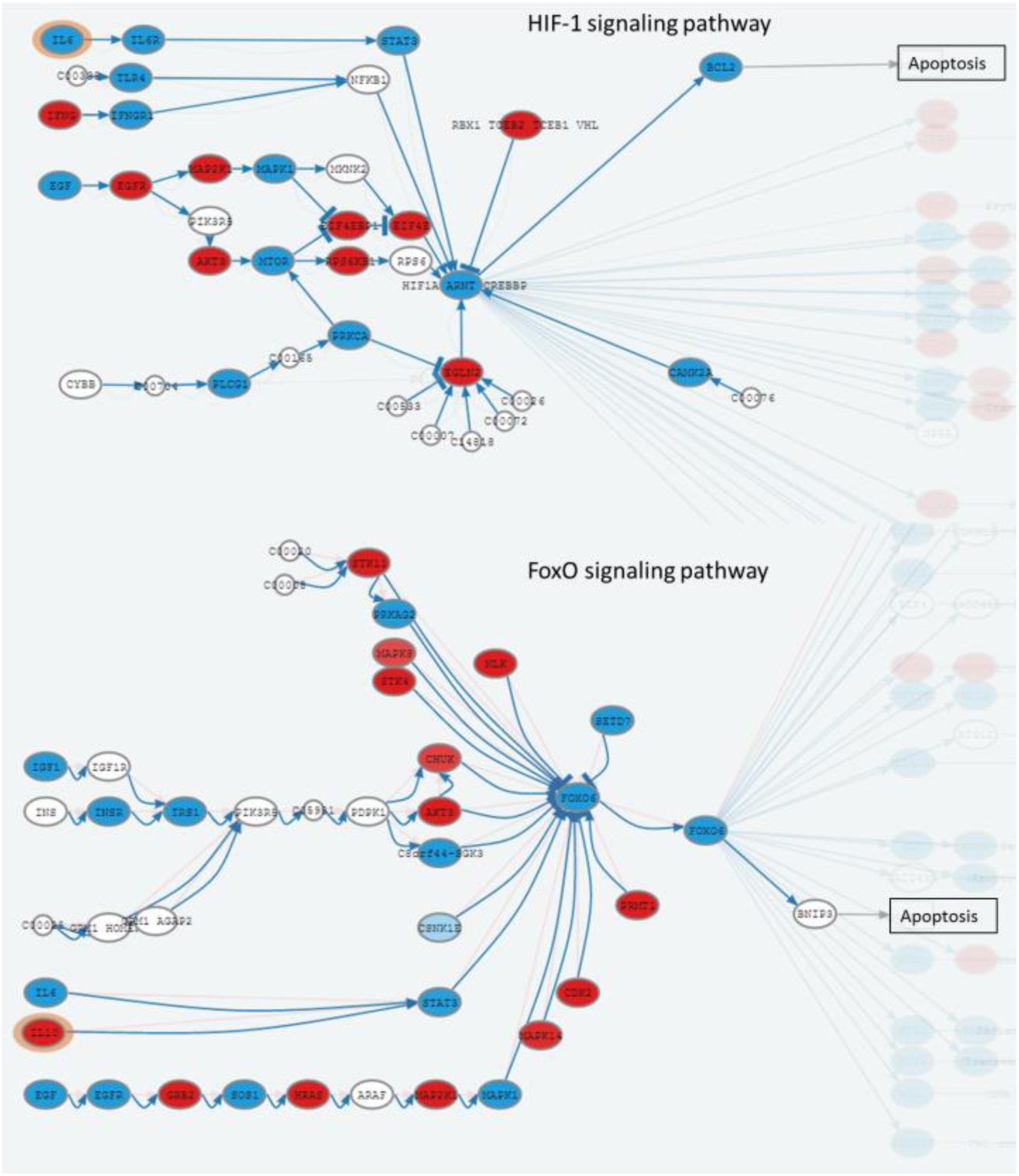
*Effector circuits* containing *Signaling circuits IL6-BCL2* in the *HIF-1 signaling pathway* and *IL10-BNIP3* in the *FoxO signaling pathway*, both highlighted in the figure. Both *circuits* contain the protein *STAT3*, known to mediate apoptosis inhibition in breast cancer.

### 3.3 The classification algorithm suggests additional prognostic genes that do not code for signaling proteins

In order to find genes that could be relevant for patient survival that are not in the signal pathways, we build a profile by combining signaling circuit activity profiles and gene expression profiles corresponding to genes outside signaling pathways (the *other-genes* profile), denoted by *path.and.other.genes.vals.* A feature selection procedure in breast cancer prognosis based on such a profile can select signaling circuits along with genes unrelated to signaling, whose activity is related to patient survival. Thus, Random Forest was again used to assess feature importance when fit with the *path.and.other.genes.vals* profile to classify survival outcome. Table 8 lists the top 5 most important gene features (the *other-genes* part of the *path.and.other.genes.vals* composed profile). These genes are of particular interest given that they might represent relevant cancer processes not included in cell signaling. Notably, the gene *ABCB5* belongs to the ATP-binding cassette subfamily B which is well known to be involved in multiple drug resistance in cancer therapy (Dean, et al., 2001), probably because its functionality of efflux transmembrane transporter. It has also been reported that *ABCB5* could mediate cell-to-cell fusion and contribute to breast cancer chemoresistance in expressing breast tumors (Frank, et al., 2005; Frank, et al., 2003). In addition, *ABCB5*, as a “pro-survival” gene, has been suggested to be a potential target against drug resistant breast cancer cells (Yang, et al., 2010). Besides, ABCB5 has been linked to melanoma (Wilson, et al., 2014). *LMO4* encodes a LIM-domain protein that has been reported as an essential mediator of cell cycle progression in ErbB2/HER2/Neu-induced breast cancer which is characterized by poor survival due to high proliferation and metastasis rates (Matthews, et al., 2013; Montañez-Wiscovich, et al., 2009). It has been reported that *LMO4* interacts with the renowned tumor suppressor *BRCA1* and inhibits *BRCA1* activity (Sum, et al., 2002; Sutherland, et al., 2003). *OPA1* encodes a mitochondrial fusion protein which might be a target for mitochondrial apoptotic effectors (Olichon, et al., 2003), such as sorafenib (Zhao, et al., 2013). The role in cancer survival played by two most important genes according to the predictor, *VPS72* and *CHADL,* is not as clear from the literature. It is worth mentioning that a mutation in VPS72 in cervix cancer with a high FATHMM pathogenicity score (Shihab, et al., 2015) is described in the COSMIC database (entry COSM458603). Regarding *CHADL*, it has been related to chondrocyte differentiation (Tillgren, et al., 2015) and extracellular matrix remodeling (Barallobre-Barreiro, et al., 2012). Therefore, both genes are potentially involved in cancer processes, which suggest that further investigation of the complete list of top-ranked *other-genes* could render new cancer drivers and potential therapeutic targets. An expanded list containing the top 50 most important features among the *other-genes* can be found in Supplementary Table 4, in which many genes with cancer-related functions can be seen. Functions for the genes have been taken from their Uniprot (UniProt_Consortium, 2015) annotations and, when absent, from GeneCards annotations (Stelzer, et al., 2016).

**Table 8.**
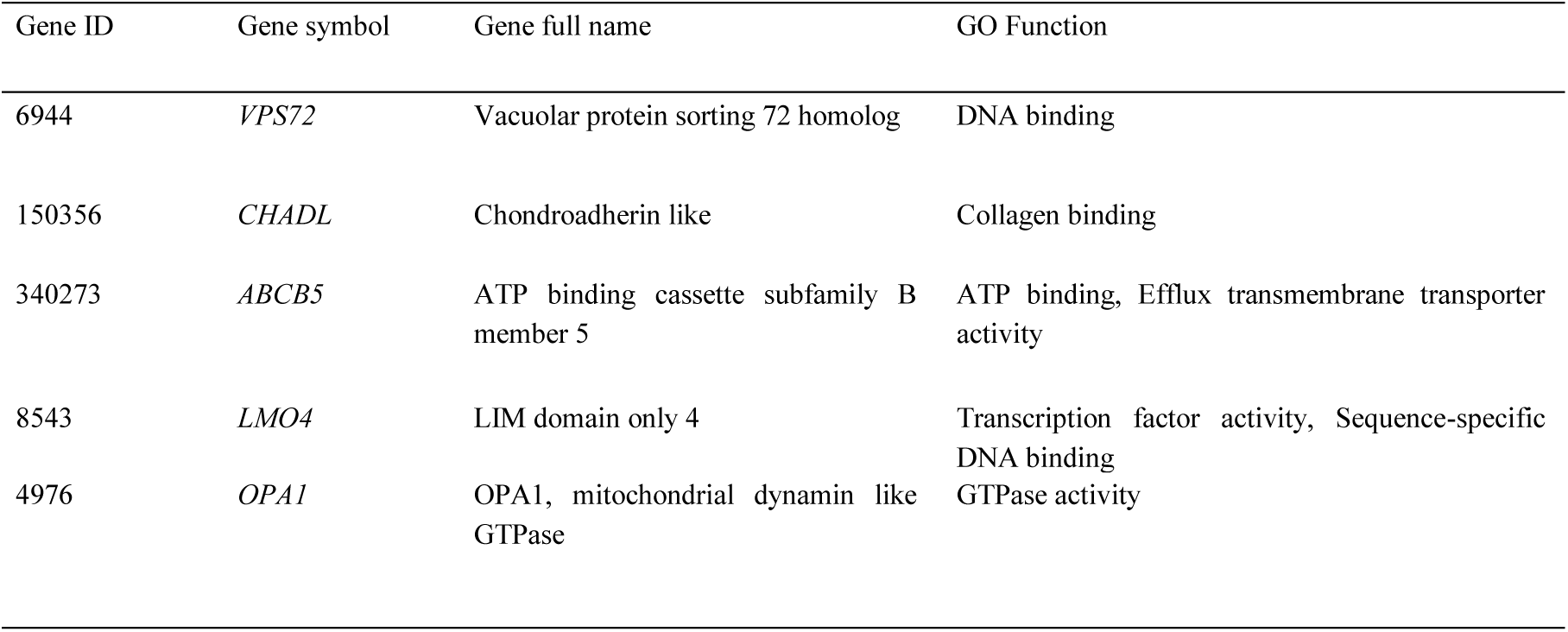
Top five genes unrelated to signaling ranked by importance in the classification of survival outcome by fitting Random Forests with *path.and.other.genes.vals* profile, along their functions as annotated in Gene Ontology.

### 3.4 Availability of data and results

All experiments are produced with R and codes are available via https://github.com/YunlongJiao/hipathiaCancerPrognosis.

There is an R package available at https://github.com/babelomics/hipathia. Additionally, there is a web interface to the hipathia methodology that includes prediction functionalities, which is freely available at: http://hipathia.babelomics.org/.

## Conclusions

In this study we have proposed a novel scheme to classify survival outcome for breast cancer patients based on mechanistic features consisting of signaling pathway activity profiles. We applied a pathway activity analysis method (Hidalgo, et al., 2017) to recode gene expression profiles into activity values of signaling circuits and demonstrated that, making use of the state-of-the-art computational tools, signaling circuit activity yields better prediction in breast cancer prognosis than gene expression. An additional advantage is that the identified pathway-level biomarkers are mechanistic signatures whose contribution to cancer progression can be readily interpreted in terms of the underlying cellular functions and biological processes.

The three feature sets *path.genes.vals, eff.vals* and *path.vals* are composed by the same genes (those present in the pathways). However, the prediction performance of the genes recoded into circuits activity values with the hipathia method (*eff.vals* and *path.vals*) clearly outperforms (see Table 4) to those of the original genes (*path.genes.vals*). Moreover, predictors based on circuits (*eff.vals* and *path.vals*) have similar performance (see Table 4) to predictors based on all the genes (*genes.vals*), which include more information than the subset of genes. It is worth noting that genes in the circuits represent only 12% of the total number of genes, but have the same predictive performance, which suggests that combining the genes into circuits provides a real added value for prediction purposes.

Although a significant improvement of the performance was not observed when the expression values of *other-genes* were concatenated to the activity values of signaling circuits, the analysis based on the combination of both data provides an interesting perspective regarding the interpretation of the biomarkers detected. In fact, the selected genes from the category of *other-genes* represent other aspects of the mechanism of the disease not explained by cell signaling. This approach allows expanding the scope of the analysis beyond the processes included in the pathways modeled.

Central to our proposed scheme is the idea of promoting gene-level analysis to pathway-level analysis by obtaining patient-specific personalized profiles of signaling circuit activity. Reliable models of pathway activity (Hidalgo, et al., 2017) can be used to derive robust multigenic biomarkers, similar to the popular MammaPrint (van′t Veer and Bernards, 2008), which in addition account properly for the underlying disease mechanisms or mechanisms of drug action.

## Funding

This work was supported by the European Union 7th Framework Program through the Marie Curie ITN MLPM grant No 316861, by the European Research Council grant ERC-SMAC-280032, by grants BIO2014-57291-R from the Spanish Ministry of Economy and Competitiveness and “Plataforma de Recursos Biomoleculares y Bioinformáticos” PT13/0001/0007 from the ISCIII, both co-funded with European Regional Development Funds (ERDF); and EU H2020-INFRADEV-1-2015-1 ELIXIR-EXCELERATE (ref. 676559)

